# Mutation-Profile-Based Methods for Understanding Selection Forces in Cancer Somatic Mutations: A Comparative Analysis

**DOI:** 10.1101/021147

**Authors:** Zhan Zhou, Yangyun Zou, Gangbiao Liu, Jingqi Zhou, Jingcheng Wu, Shimin Zhao, Zhixi Su, Xun Gu

## Abstract

Human genes exhibit different effects on fitness in cancer and normal cells. Here, we present an evolutionary approach to measure the selection pressure on human genes, using the well-known ratio of the nonsynonymous to synonymous substitution rate in both cancer genomes (*C_N_/C_S_*) and normal populations (*p_N_/p_S_*). A new mutation-profile-based method that adopts sample-specific mutation rate profiles instead of conventional substitution models was developed. We found that cancer-specific selection pressure is quite different from the selection pressure at the species and population levels. Both the relaxation of purifying selection on passenger mutations and the positive selection of driver mutations may contribute to the increased *C_N_/C_S_* values of human genes in cancer genomes compared with the *p_N_/p_S_* values in human populations. The *C_N_/C_S_* values also contribute to the improved classification of cancer genes and a better understanding of the onco-functionalization of cancer genes during oncogenesis. The use of our computational pipeline to identify cancer-specific positively and negatively selected genes may provide useful information for understanding the evolution of cancers and identifying possible targets for therapeutic intervention.

## INTRODUCTION

Since the pioneering work of Cairns and Nowell [1, 2], the evolutionary concept of cancer progression has been widely accepted [3-7]. In this model, cancer cells evolve through random somatic mutations and epigenetic changes that may alter several crucial pathways, a process that is followed by clonal selection of the resulting cells. Consequently, cancer cells can survive and proliferate under deleterious circumstances [8, 9]. Therefore, knowledge of evolutionary dynamics will benefit our understanding of cancer initiation and progression. For example, there are two types of somatic mutations in cancer genomes: driver mutations and passenger mutations [10, 11]. Driver mutations are those that confer a selective advantage on cancer cells, as indicated by statistical evidence of positive selection. Passenger mutations do not confer a clonal growth advantage and are usually considered neutral in cancer. However, some passenger mutations in protein-coding regions that would have potentially deleterious effects on cancer cells may be under negative selection in cancer [12, 13].

Cancer somatic mutations, especially driver mutations, promote the cancer specific functionalization of cancer-associated genes, i.e., onco-functionalization. Onco-57 functionalization of cancer-associated genes would promote cancer initiation and progression. For example, oncogenes may gain new functions during carcinogenesis, which could be considered cancer-specific neo-functionalization [14]. By contrast, the mutation of tumor suppressor genes to cause a loss or reduction of their function could be considered cancer-specific non-functionalization [15].

Analyses of large-scale cancer somatic mutation data have revealed that the effects of positive selection are much stronger on cancer cells than on germline cells [16, 17]. Given that many of the positively selected genes in tumor development act as the driving force behind tumor initiation and development and are thus considered “driver genes”, it is understandable that almost all previous studies have focused on positively selected genes in cancer genomes [3, 18-21]. Nevertheless, we have realized that an alternative approach, i.e., identifying cancer-constrained genes that are highly conserved in tumor cell populations (under purifying selection), is also valuable. For example, TP73, a homolog of TP53, is rarely mutated but frequently overexpressed in tumor cells. TP73 has been reported to activate the expression of glucose-6-phosphate dehydrogenase and support the proliferation of human cancer cells [22]. As essential genes are crucial for carcinogenesis, progression and metastasis, this idea may be advantageous in addressing issues related to drug resistance in cancer therapies, especially in cancers with high intratumor heterogeneity.

Many previous studies have used the ratio of nonsynonymous to synonymous substitution rates to identify genes that might be under strong positive selection both in organismal evolution and carcinogenesis [11, 16, 17, 23-26]. However, most of these studies applied conventional methods, which are usually based on simple nucleotide mutation/substitution models, e.g., the simplest equal-rate model assuming that every mutation or substitution pattern has the same probability [27]. Unfortunately, this may not be a realistic biological model because many recent cancer genomics studies have shown that mutation profiles vary greatly between different cancer samples [17, 28]. In addition, context-dependent mutation bias (i.e., base-substitution profiles that are influenced by the flanking 5’ and 3’ bases of each mutated base) should also be considered [28, 29].

In this study, we describe a mutation-profile-based method to estimate the selective constraint for each gene in pan-cancer samples and human populations. In brief, the new method discards an unrealistic assumption inherent in the equal-rate model that every mutation or substitution pattern has the same probability [27]. This assumption can lead to nontrivial biased estimations when it is significantly violated. By contrast, our method implements an empirical nucleotide mutation model that simultaneously considers account several factors, including single-base mutation patterns, local-94 specific effects of surrounding DNA regions, and tissue/cancer types. Using simple somatic mutations from 9,155 tumor-normal paired whole-exome/genome sequences (ICGC Release 20), as well as rare germline substitutions from 6,500 exome sequences from the National Heart, Lung, and Blood Institute (NHLBI) Grant Opportunity (GO) Exome Sequencing Project (ESP), as references, we used this mutation-profile-based method to identify selective constraints on human genes, especially cancer-associated genes, in cancer cells. Our results may provide useful information for the precise classification of known cancer-associated genes and for an improved understanding of the evolution of cancers.

## RESULTS

### The mutation rate profiles in cancer genomes and human populations differ

Estimating evolutionary selective pressure on human genes is a practical method for inferring the functional importance of genes in a specific population. By comparing selective pressures, we may be able to identify different functional and fitness effects of human genes in cancer and normal cells. The conventional method for measuring selective pressure is to calculate the ratio of nonsynonymous to synonymous substitution rates using the equal-rate method [27], which assumes equal substitution rates among different nucleotides. In this study, we used the cancer somatic mutations from 9,155 tumor-normal pairs from ICGC (Release 20) as well as rare variants (minor allele frequency <0.01%) from 6,500 exome sequences from ESP as a reference. We used these data to compare the empirical mutation rate profiles of cancer somatic mutations and germline substitutions using 96 substitution classifications [28, 29]. The empirical mutation rate profiles reveal the prevalence of each substitution pattern for point mutations and present not only the substitution types but also the sequence context (see Methods). The exonic mutation profiles of cancer somatic mutations and germline substitutions are both enriched in C-to-T transitions (Figure 1). The mutation rates for each trinucleotide context differ from each other, and the ratio of transition to transversion for each trinucleotide context is much greater than 1:2 for both cancer somatic mutations (ratio=2.70±0.47) and germline rare variants (ratio=3.28±0.53) (Supplementary Figure S1). These different mutation profiles may lead to different biological progressions in carcinogenesis, as depicted in several publications [19, 28]. For example, the mutation profiles of melanoma are highly enriched in C-to-T transitions, indicating a direct mutagenic role of ultraviolet (UV) light in melanoma pathogenesis [30]. Thus, it is inappropriate to use conventional methods such as the equal-rate model to measure selective pressure because this approach ignores the mutation bias of different nucleotide substitution types.

**Figure 1.**
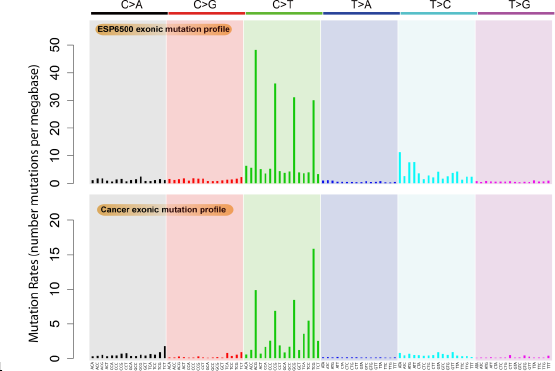
Mutation profiles of cancer somatic mutations and germline substitutions, including the exonic mutation profile of 9,155 cancer samples and the exonic mutation profile of ESP6500.

### Measuring selective pressure on human genes in cancer and germline cells using the mutation-profile-based method

We therefore formulated an evolutionary approach that was designed specifically to estimate the selective pressure imposed on human genes in cancer cells and then identify genes that had undergone positive and purifying selection in cancer cells compared with in normal cells (see Figure 2 for an illustration). In cancer genomics, distinguishing synonymous from nonsynonymous somatic mutations is straightforward. We developed the mutation-profile-based method to estimate the *C_N_/C_S_* ratio of each human gene based on the mutation profiles of cancer somatic mutations and the *p_N_/p_S_* ratio for germline substitutions. In contrast to the equal-rate method [27], our method considers differences in substitution rates and uses the overall mutation rate profile as the weight matrix (Figure 1).

**Figure 2.**
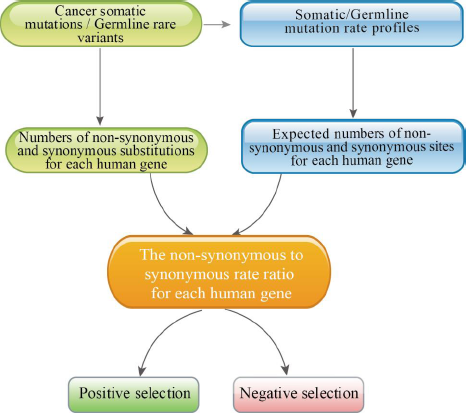
The pipeline used to identify positively and negatively selected cancer genes with the mutation-profile-based method.

We calculated the expected number of nonsynonymous and synonymous sites based on the exonic mutation rate profiles. We then counted the number of nonsynonymous and synonymous substitutions in the protein-coding region of each human gene for all cancer somatic mutations or germline substitutions. A χ^2^ test was performed to identify the genes whose *C_N_/C_S_* values were either significantly greater than one or less than one, which indicates positive or negative (purifying) selection, respectively. Of the 16,953 genes with at least one germline substitution and cancer somatic mutation, the overall *C_N_/C_S_* value for cancer somatic mutations (mean±s.e.=1.199±0.008) was much greater than the overall *p_N_/p_S_* of germline substitutions (mean±s.e.=0.738±0.005) (Wilcoxon test, p<2.2×10^−16^) (Table 1A, Supplementary Table S1). In the cancer genomes, 365 genes had *C_N_/C_S_* values 155 significantly greater than one, and 923 genes had *C_N_/C_S_* values significantly less than one (χ^2^ test, p<0.01, FDR<0.1). By contrast, germline substitutions included only 24 genes with *p_N_/p_S_* values significantly greater than one, whereas 4,897 genes had *p_N_/p_S_* values significantly less than one (χ^2^ test, p<0.01, FDR<0.1). Of these 365 cancer positively selected genes, only one gene (*RSRC1*) also exhibited positive selection whereas 117 genes exhibited negative selection in germline substitutions. Additionally, 500 cancer negatively selected genes did not exhibit significant negative selection in germline substitutions. These genes may therefore be under different selective pressure in cancer and germline genomes.

**Table 1.**
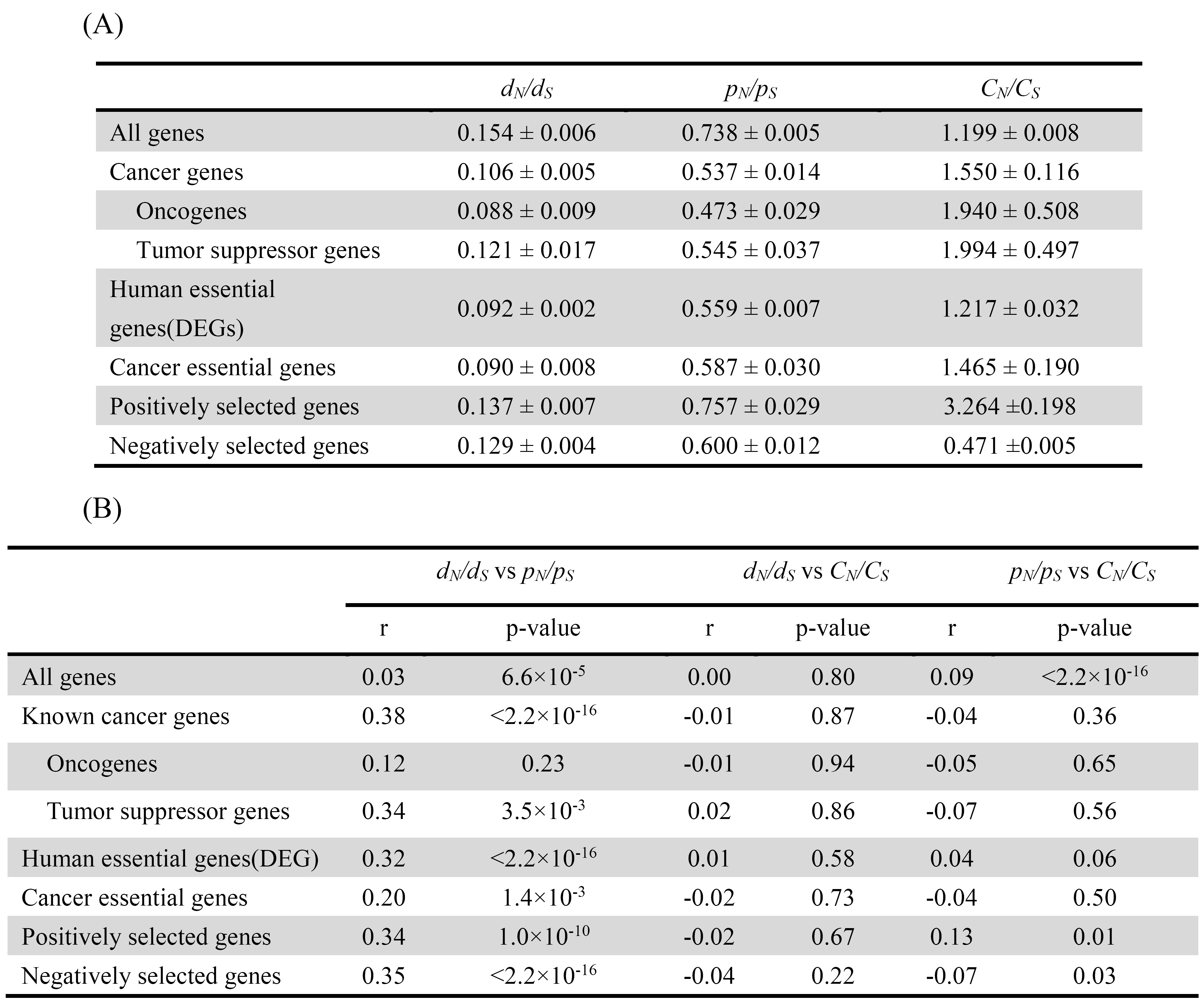
The *ω* ratios (*d_N_/d_S_*, *p_N_/p_S_*, *C_N_/C_S_* values) (A) and the correlations of the *ω* 654 ratios (B) for the different gene sets for the human-mouse orthologs and for germline and cancer somatic mutations. The positively and negatively selected genes indicates the genes that are under positive and negative selection in cancer cells, respectively (χ^2^ test, p<0.01, FDR<0.1).

Previous studies have attributed elevated *C_N_/C_S_* values to the relaxation of purifying selection [16] or increased positive selection of globally expressed genes [17]. Our results show that the number of genes under positive selection increased, whereas the number of genes under negative selection decreased, in cancer genomes compared with germline genomes. This result indicates that both the relaxation of purifying selection on passenger mutations and the positive selection of driver mutations may contribute to the increased *C_N_/C_S_* values of human genes in cancer genomes.

### Selection pressures on cancer-associated genes

The Cancer Gene Census (CGC) [31, 32] contains more than 500 cancer-associated genes that have been reported in the literature to exhibit mutations and that are causally implicated in cancer development. Of those genes, 553 were included in the 16,953 genes that we tested. These known cancer genes have significantly greater *C_N_/C_S_* values (Wilcoxon test, p=2.9×10^−10^) for cancer somatic mutations but significantly lower *p_N_/p_S_* values for germline substitutions (Wilcoxon test, p<2.2×10^−16^) than other genes (Table 1A). For selection over longer evolutionary time scales, we extracted the *d_N_/d_S_* values between human-mouse orthologs from the Ensembl database (Release 75) [33]. The known cancer genes have significantly lower human-mouse *d_N_/d_S_* values than other human genes (Wilcoxon test, p<2.2×10^−16^). These results support the work of Thomas *et al.* [34], who showed that known cancer genes may be more constrained and more important than other genes at the species and population levels, especially for oncogenes. By contrast, known cancer genes are more likely to gain onco-functional somatic mutations in cancer than other genes.

Among the 365 cancer positively selected genes, 45 (12.3%) genes are known cancer genes, indicating that cancer genes are significantly enriched in cancer positively selected genes (Fisher’s Exact Test, p=6.7×10^−15^). When we choose a more stringent cut-off of p<10^−5^, 17 of the 29 (58.6%) positively selected genes are known cancer genes, according to the CGC, and the work of Lawrence *et al.* [20] and Kandoth *et al.* [35], such as the well-known cancer drivers *TP53*, *KRAS*, *PIK3CA*, and *BRAF.* (Supplementary Table S2). In addition, the 29 strong positively selected genes are significantly enriched in biological processes related to cancer, according to the functional analysis using DAVID v6.7 [36] (Supplementary Table S3). Some cancer genes also show negative selection in cancer genomes, such as the oncogene *MLLT3* (*C_N_/C_S_*=0.11, p=3.14×10^−44^, FDR=5.52×10^−41^). The *MLL-MLLT3* gene fusion is the main mutation type of *MLLT3* that drives tumorigenesis in acute leukemia [37]. Interestingly, *MLLT3* has recurrent synonymous mutations at amino acid positions 200 to 168 (S166S, 8/9155; S167S, 33/9155; S168S, 23/9155).

Using the *C_N_/C_S_* values, we classified known cancer genes according to the selection pressure on these genes in cancer cells, as well as their onco-functionalization in oncogenesis (Table 2). The most important two classes are oncogenes and tumor suppressor genes that are under strong positive selection, such as *TP53*, the most famous tumor suppressor gene [38], which shows strong positive selection pressure (*C_N_/C_S_*=32.57, p=1.06×10^−159^, FDR=6.55×10^−156^). The non-synonymous mutations of *TP53* with onco-nonfunctionalization are distributed in a wide range of cancers. The oncogene *KRAS* [39] also showed a strong positive selection pressure (*C_N_/C_S_*=45.88, p=4.25×10^−87^, FDR=1.74×10^−83^). Recurrent non-synonymous mutation with onco-210 neofunctionalization of *KRAS* are highly enriched in codons 12 and 13; mutations in these codons represent 79.4% and 8.0% of all non-synonymous mutations of *KRAS*.

**Table 2.** Classification of cancer genes according to cancer-specific selection pressures

|  | #Positive selection | #Negative selection | #Neutral |
| --- | --- | --- | --- |
| Known cancer genes | 45 | 29 | 479 |
| Oncogenes | 11 | 7 | 79 |
| Tumor suppressor genes | 10 | 6 | 54 |

We also observed 12 cancer positively selected genes (p<10^−5^) that have not been reported as cancer-associated genes. These genes are recurrently mutated in several tumor types and are potential cancer driver genes. According to the mouse insertional mutagenesis experiments [40], three of these genes (*DMD, MYO9A*, and *COL5A2*) have been identified as cancer-causing genes [41-44].

When we chose a more stringent cut-off of p<10^−5^ for cancer negatively selected genes, we found 112 genes that showed an enrichment in the Notch signaling pathway (Supplementary Table S3). Forty-seven of the 112 negatively selected genes showed more stringent selective constraint in cancer cells than in normal cells (*p_N_/p_S_* > *C_N_/C_S_*, p>0.05 for *p_N_/p_S_*). It would be quite valuable to uncover the roles of these evolutionarily conserved genes in cancer cells. Out of the 47 genes, 14 genes showed a significantly increased expression level in cancers than in normal tissues (fold change>2, p<10^−4^) (Supplementary Table S4). For example, *SPRR3*, a member of the small proline-rich protein family, is under purifying selection in cancer cells (*C_N_/C_S_*=0.27, p=5.73×10^−11^, FDR=1.91×10^−8^) and neutral selection in germline cells (*p_N_/p_S_* =0.88, p=0.75, FDR=0.37). It has been reported that *SPRR3* is overexpressed in several tumor types, and is associated with tumor cell proliferation and invasion. Therefore, *SPRR3* could be a potential biomarker and novel therapeutic target [45-47].

We also examined essential genes during human development and cancer development. We extracted 2,452 human orthologs of mouse essential genes from DEG10 (the Database of Essential Genes) [48]. These genes, which are human orthologs of known essential genes in mice [49], are critical for cell survival and are therefore more conserved than other genes at the species and population levels. Here, we found that human orthologs of mouse essential genes have significantly lower *d_N_/d_S_* 236 values (measured between human-mouse orthologs) and lower *p_N_/p_S_* values for germline substitutions but similar *C_N_/C_S_* values for cancer somatic mutations compared with the values for non-essential genes (Table 1A). Human orthologs of mouse essential genes are also enriched among cancer positively selected genes. Eighteen of the twenty-240 nine (62.1%) positively selected genes (p<10^−5^) are human orthologs of mouse essential genes (Supplementary Table S2). We also used the human orthologs of mouse essential genes from OGEE (the database of Online GEne Essentiality) [50] to confirm these results (Supplementary Table S2).

Cancer essential genes were identified by performing genome-scale pooled RNAi screens. RNAi screens with the 45k shRNA pool in 12 cancer cell lines, including small-cell lung cancer, non-small-cell lung cancer, glioblastoma, chronic myelogenous leukemia, and lymphocytic leukemia, revealed 268 common essential genes [51]. Compared to other human genes, these cancer essential genes have significantly lower *d_N_/d_S_* values and lower *p_N_/p_S_* values for germline substitutions and greater *C_N_/C_S_* values for cancer somatic mutations ((Table 1A), suggesting a functional shift of these genes in human populations and cancer cells.

We further tested the correlations of the *d_N_/d_S_*, *p_N_/p_S_* and *C_N_/C_S_* values of human genes for human-mouse orthologs, germline substitutions and cancer somatic mutations to compare selective pressures among species, populations and cancer cells (Table 1B). For different gene sets, the *d_N_/d_S_* values show a weak positive correlation with the *p_N_/p_S_* values, but no significant correlation with *C_N_/C_S_* values. The *p_N_/p_S_* values and *C_N_/C_S_* values also do not have significant correlation for different gene sets. These results indicate that the cancer-specific selection pressure is quite different from the selection pressure at the species and population levels.

### Selection pressure among different cancer types

As cancer is highly heterogeneous, we further analyzed the selection pressure of human genes in different cancer types. The 9,155 tumor samples from the ICGC database could be classified as 20 cancer types according to the primary site. The overall *C_N_/C_S_* values for the cancer somatic mutations in the different cancer types ranged from 1.078±0.022 to 1.827±0.013 (mean±s.e., Table 3). The detected positively and negatively selected genes (χ^2^ test, p<0.01) varied in the different cancer types (Supplementary Table S5). Due to the limited number of tumor samples and somatic mutations for each cancer type, particularly in the cancer types with low mutation rates, our method might not be sensitive enough to detect the selection pressure for each gene. For example, only one positively selected gene was detected in bone cancer (IDH1) and nervous system cancer (ALK), respectively. There were also three genes (TP53, PIK3CA and KRAS) that showed positive selection in more than five cancer types. In particular, TP53 showed positive selection in 15 cancer types. On the other hand, more genes (164/188, 87.2%) were under positive selection in only one cancer type. We also found that six genes (TBP, EP400, DSPP, MUC21, MLLT3, and MUC2) were under negative selection in more than five cancer types. These genes also showed negative selection at the species and population levels. Furthermore, 85.8% (2,417/2,817) of genes showed negative selection in only one cancer type. These results indicate the divergence of selection pressure in different cancer types.

**Table 3.**
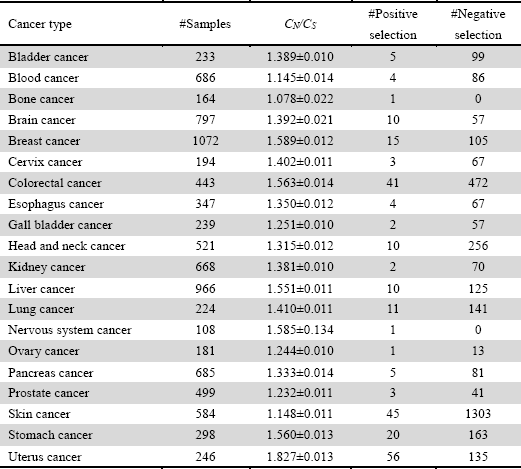
The selection pressure in different cancer types

| Cancer type | #Samples | $C_N/C_S$ | #Positive selection | #Negative selection |
| --- | --- | --- | --- | --- |
| Bladder cancer | 233 | 1.389±0.010 | 5 | 99 |
| Blood cancer | 686 | 1.145±0.014 | 4 | 86 |
| Bone cancer | 164 | 1.078±0.022 | 1 | 0 |
| Brain cancer | 797 | 1.392±0.021 | 10 | 57 |
| Breast cancer | 1072 | 1.589±0.012 | 15 | 105 |
| Cervix cancer | 194 | 1.402±0.011 | 3 | 67 |
| Colorectal cancer | 443 | 1.563±0.014 | 41 | 472 |
| Esophagus cancer | 347 | 1.350±0.012 | 4 | 67 |
| Gall bladder cancer | 239 | 1.251±0.010 | 2 | 57 |
| Head and neck cancer | 521 | 1.315±0.012 | 10 | 256 |
| Kidney cancer | 668 | 1.381±0.010 | 2 | 70 |
| Liver cancer | 966 | 1.551±0.011 | 10 | 125 |
| Lung cancer | 224 | 1.410±0.011 | 11 | 141 |
| Nervous system cancer | 108 | 1.585±0.134 | 1 | 0 |
| Ovary cancer | 181 | 1.244±0.010 | 1 | 13 |
| Pancreas cancer | 685 | 1.333±0.014 | 5 | 81 |
| Prostate cancer | 499 | 1.232±0.011 | 3 | 41 |
| Skin cancer | 584 | 1.148±0.011 | 45 | 1303 |
| Stomach cancer | 298 | 1.560±0.013 | 20 | 163 |
| Uterus cancer | 246 | 1.827±0.013 | 56 | 135 |

### Comparison of the equal-rate model and empirical mutation profile model

Considering that different nucleotide substitution models might provide varying estimates, we used the equal-rate method [27] as the simplest model to calculate the expected numbers of nonsynonymous and synonymous sites. The overall *C_N_/C_S_* value for cancer somatic mutations (mean±s.e.=0.892±0.006) is greater than the *p_N_/p_S_* value for germline substitutions (mean±s.e.=0.633±0.004) for the 16,953 genes (Supplementary Table S1) but lower than that calculated using the mutation-profile-289 based method (Wilcoxon test, p<2.2×10^−16^) (Figure 3A). Consequently, the number of genes with *C_N_/C_S_* values >1 (χ^2^ test, p<0.01, FDR<0.1) is much lower than those calculated using the exonic mutation profiles (37 versus 365), whereas the number of genes with *C_N_/C_S_* values <1 (χ^2^ test, p<0.01, FDR<0.1) is much greater (2851 versus 923) (Figure 3B and 3C).

**Figure 3.**
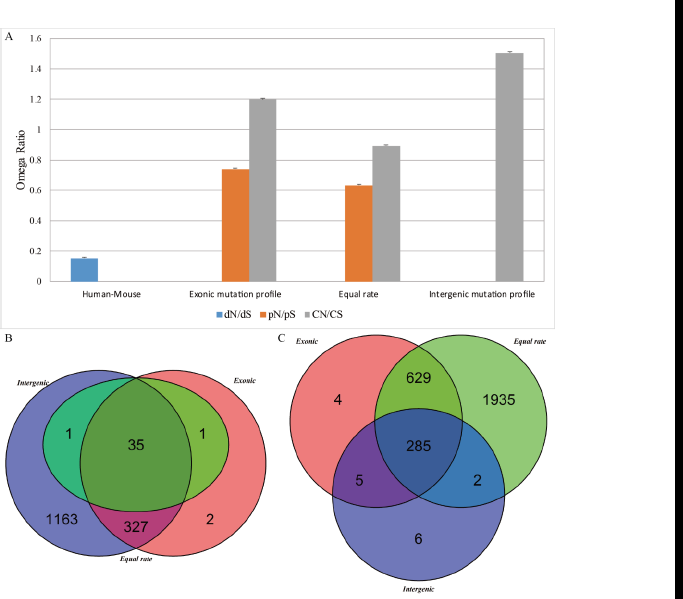
The overall omega ratio (A) and overlap of cancer positively selected (B) and negatively selected (C) genes based on different models.

We also used the intergenic mutation rate profile from 2,900 tumor-normal whole genome sequences, which are included in the 9,155 cancer samples of ICGC database, to calculate the *C_N_/C_S_* value for cancer somatic mutations. The overall *C_N_/C_S_* value (mean±s.e.=1.503±0.010) is greater than that calculated from the exonic mutation rate profile (mean±s.e.=1.199±0.008) (Wilcoxon test, p<2.2×10^−16^), resulting in more positively selected genes (1526 versus 365) and fewer negatively selected genes (298 versus 923) (Figure 3B and 3C).

The equal-rate method ignores the mutation rate bias between different substitution types, especially the ratio of transition to transversion, leading to underestimation of the *C_N_/C_S_* ratio. Therefore, the equal-rate method is strict for positive selection detection but relaxed for the detection of negative selection [52]. In contrast, the mutation-profile-based method considers the mutation bias, which can be depicted as the internal variance between mutation rates of different substitution types. Thus, the mutation-307 profile-based method can correct the underestimation of the *C_N_/C_S_* ratio estimated by the equal-rate method. Furthermore, the mutation-profile-based method would also increase the false-positive results for detecting positively selected genes but be more conservative in detecting negatively selected genes. The mutation bias may simulate the detection of genes under strong selection pressure but may suppress the detection of genes under weak selection pressure.

## DISCUSSION

A key goal of cancer research is to identify cancer-associated genes, such as oncogenes and tumor suppressor genes, that might promote tumor occurrence and progression when mutated [28]. Instead of searching for cancer-causing genes with multiple driver mutations, an alternative approach is to identify cancer essential genes in tumor cell populations because they are crucial for carcinogenesis, progression and metastasis. Cancer essential genes are important for the growth and survival of cancer cells [51] and are expected to be highly conserved in cancer cells. In this study, we aimed to detect both cancer-specific positively and negatively selected genes using a molecular evolution approach.

Based on analyses of large-scale cancer somatic mutation data derived from The Cancer Genome Atlas (TCGA) or International Cancer Genome Consortium (ICGC), previous studies identified important differences between the evolutionary dynamics of cancer somatic cells and whole organisms [6, 16, 18]. However, these studies applied canonical nucleotide substitution models to identify the molecular signatures of natural selection in cancer cells or human populations and neglected the apparently different mutation profiles of these cell types. Here, we developed a new mutation-profile-based method to calculate the *C_N_/C_S_* values of human genes for cancer somatic mutations. In our results, a large number of known cancer genes did not show significant positive selection according to our analysis. One possible reason for this finding suggests that positive selection for driver mutations is obscured by the relaxed purifying selection of passenger mutations. Additionally, among the strong positively selected genes, more than half are known cancer genes. Another possible reason might be that the main mutation type of more than 300 cancer-associated genes is translocation or copy number variation, rather than point mutation. Furthermore, some of the positively selected genes might also be related to cancer, such as *DMD*, *MYO9A*, and *COL5A2*, which have been identified as cancer-causing genes based on mouse insertional mutagenesis experiments [40].

Two prerequisites are crucial to properly apply the mutation-profile-based method. First, a large number of samples with similar mutation profiles are necessary to increase the power of selection pressure detection. Second, a subset of nucleotide substitutions should be chosen to represent the background neutral mutation profiles of the samples. In this study, because of the limited number of cancer samples, especially the number of whole-genome sequenced tumor-normal tissue pairs, we pooled all samples to analyze pan-cancer-level selection pressures. However, cancer somatic mutation profiles are well known to be heterogeneous among different cancer types, even for samples with the same tissue origin [19, 20, 28, 35]. As the number of sequenced cancer genomes increases, we will be able to classify cancer samples by their specific mutation profiles and infer evolutionarily selective pressures more precisely using the mutation-353 profile-based method.

Background neutral mutation profiles can be calculated based on intergenic regions from the corresponding samples. In this study, we assumed that most of the exonic somatic mutations in the cancer samples do not have significant effects on the fitness of cancer cells. Under this assumption, we can apply the mutation profiles of coding regions to approximate the background. The exonic mutation profiles used in our mutation-profile-based method considered the weight of the 96 substitution classifications within the cancer exomes, which may reflect the mutation bias of different substitution types within the protein-coding regions. This method would correct the underestimation of the *C_N_/C_S_* value that occurs with the equal-rate method [52]. The mutation-profile-based method is more sensitive for the detection of positive selection but more conservative for the detection of negative selection compared with the equal-rate method. As more tumor-normal whole genome sequence data become available, it would be better to choose suitable mutation profiles for the mutation-367 profile-based method. With the expansion of these data in the future, we may apply more precise methods to identify neutral background mutation properties.

## MATERIALS AND METHODS

### Datasets

Cancer somatic mutation data from 9,155 cancer samples corresponding to 20 373 primary sites were extracted from the ICGC Data Portal (http://dcc.icgc.org, Release 20), which includes 36,985,985 somatic mutations and small insertions/deletions. Data on rare human protein-coding variants (minor allele frequency <0.01%) from 6,500 human exomes (ESP6500) were extracted from the NHLBI GO Exome Sequencing Project (http://evs.gs.washington.edu/EVS, Exome Variant Server NGESPE, Seattle, WA). A total of 572 known cancer genes were extracted from the Cancer Gene Census (http://cancer.sanger.ac.uk/cancergenome/projects/census/, COSMIC v72) [31, 32].

Human gene sequences and annotations were extracted from the Ensembl database (GRCh37, Release 75) [33]. For each gene, only the longest transcript was selected for the subsequent analyses. The *d_N_/d_S_* values between human-mouse orthologs were extracted from the Ensembl database. The HGNC (HUGO Gene Nomenclature Committee) database [53] (http://www.genenames.org/) and the Genecards database [54] (http://www.genecards.org) were used to map the gene IDs from different datasets. DAVID v6.7 was utilized for the functional annotation analysis [36]. ANNOVAR was utilized to perform biological and functional annotations of the cancer somatic mutations and germline substitutions [55]. The Oncomine database [56] (https://www.oncomine.org) was used to compare the gene expression level of negatively selected genes between cancer and normal tissues. The human orthologs of mouse essential genes were extracted from the DEG 10 [48] and the OGEE v2 [50] 392 databases.

### Statistical measure for gene-specific selection pressure in cancer evolution (*C_N_/C_S_*)

In cancer genomics, distinguishing synonymous from nonsynonymous somatic mutations is straightforward. Thus, given a set of independent cancer samples, the ratio of nonsynonymous counts (N) to synonymous counts (S) of a gene, denoted by *N/S*, is simply given by the sum over all samples, under the assumption of no double mutations at the same nucleotide site (e.g., for the observed mutation A>C, the mutation path A>G>C is almost impossible in cancers). To further explore the underlying mechanisms, the *N/S* ratio must be normalized by *L_N_/L_S_*, that is,

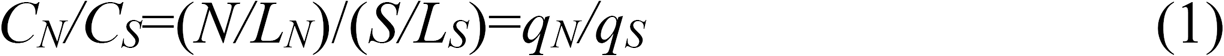

where *L_N_* is the number of expected nonsynonymous sites and *L_S_* is the number of expected synonymous sites. Note that *C_N_/C_S_* is specific for cancer somatic mutations, to avoid notation confusions with *d_N_/d_S_* in molecular evolution and *p_N_/p_S_* in population genetics. To avoid a calculation error for the small sample size, 0.5 was added to each parameter for the calculation of *C_N_/C_S_* if N or S was equal to zero.

The calculation of *L_N_* and *L_S_* from the nucleotide sequence is not a trivial task. For instance, in the codon TTT (coding for amino acid Phe), the first two positions are counted as nonsynonymous sites because no synonymous changes can occur at these positions. At the third position, the transition change (T>C) is synonymous, whereas the remaining two transversion changes (T>A and T>G) are nonsynonymous. Apparently, the weight of the third position of codon TTT as synonymous (*w_S_*) or nonsynonymous (*w_N_*) depends on the pattern of somatic mutations. At one extreme, if the transition mutation is dominant, this position should nearly be counted as a synonymous site (*w_S_*=1); at the other extreme (transversion dominant), this position would be counted as a nonsynonymous site (*w_S_*=0).

### Equal-rate model

The weight of a nucleotide as synonymous (*w_S_*) is simple when the rate of base change is the same. Let *I_S_* be the number of possible synonymous changes at a site. This is counted as *w_S_*=*I_S_*/3 synonymous and (1-*I_S_*/3) nonsynonymous. For instance, in the codon TTT (Phe), the first two positions are counted as nonsynonymous sites because no synonymous changes can occur at these positions (*w_S_*=0). The third position of codon TTT is then counted as one third of a synonymous site (*w_S_*=1/3) and two-thirds of a nonsynonymous site (*w_N_*=2/3) because only one of the three possible changes is synonymous. It is then straightforward to calculate the numbers of synonymous and nonsynonymous sites.

### Empirical mutation profile model

Substantial evidence has demonstrated that the rate of somatic mutations in cancer depends on not only the nucleotide site (e.g., synonymous or nonsynonymous sites) and the mutation type (e.g., transition or transversion) but also on the sequence context of each mutated site, i.e., the effects of near-by nucleotides on somatic mutations are nontrivial. Recent studies [28, 29, 57] proposed an empirical mutation profile of any position with base P, considering two immediate neighbor nucleotides (x, y) of a trinucleotide string denoted by xPy. Since base P has six base-change patterns (under Watson-Crick pairing) and both x and y have four types of bases, there are a total of 4×6×4=96 substitution classifications, with the empirical profile denoted by *M*(*xPy*>*xP_i_*y*), where *P_i_\** (*i*=1,2,3) for the other three bases instead of P. To determine the probability of the mutation type (*xPy*>*xP_i_*y*), we divided the number of mutations in that trinucleotide context (*xPy*>*xP_i_*y*) by the number of occurrences of the trinucleotide (*xPy*). Our computational pipeline is illustrated by the following example.

In the encoding sequence with two codons … TT*T*-ATG…., we consider the third position of codon TTT (Phe). Under the trinucleotide TTA for the mutation profile (not the codon), the corresponding three substitution configurations are given by *M*(TTA>TCA), *M*(TTA>TAA) and *M*(TTA>TGA), respectively, and the number of occurrences of TTA is *M*(TTA). Next, we consider codon TTT. Because TTT and TTC are synonymous codons but TTA and TTG are not, the probabilities that this site will be synonymous and nonsynonymous are simply given by the following:

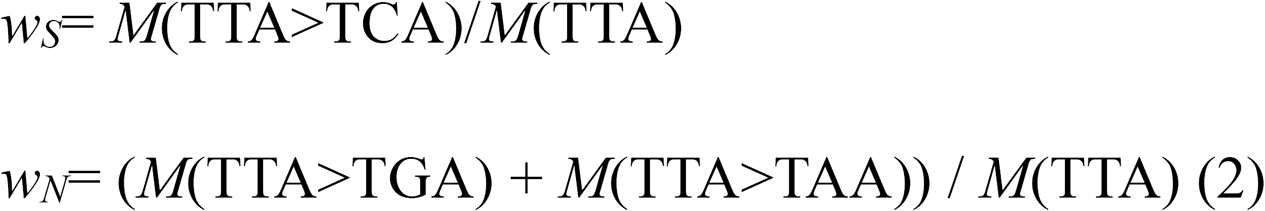

We counted all somatic mutations in the protein-coding regions of the 9,155 tumor-normal paired cancer samples, as well as all the rare protein-coding variants of the ESP6500 dataset. The mutation profiles were depicted as the mutation rate of each mutation type according to the 96 substitution classifications.

The ratio of transition to transversion for each trinucleotide context was calculated based on the mutation rate of transitions and transversions. For example, the ratio of transition to transversion for ACA=M(ACA>ATA)/(M(ACA>AAA)+M(ACA>AGA)).

### Detection of positive and negative selections

The χ^2^ test was used to compare the number of nonsynonymous and synonymous substitutions to the number of nonsynonymous and synonymous sites for each gene to test the statistical significance of the difference between the *C_N_/C_S_* values and one. Genes with *C_N_/C_S_* values significantly greater than one were classified as under positive selection in tumors, whereas genes with *C_N_/C_S_* values significantly less than one were classified as under negative, or purifying, selection. The false-discovery rate was estimated using the qvalue package from Bioconductor [58]. The software tool R was used for statistical analysis (http://www.r-project.org/).

## ACKNOWLEDGEMENTS

We are grateful to Xiaopu Wang for his help with the manuscript preparation. We would like to thank the NHLBI GO Exome Sequencing Project and its ongoing studies which produced and provided the exome variant calls for comparison: the Lung GO Sequencing Project (HL-102923), the WHI Sequencing Project (HL-102924), the Broad GO Sequencing Project (HL-102925), the Seattle GO Sequencing Project (HL-477 102926) and the Heart GO Sequencing Project (HL-103010). We also gratefully acknowledge the clinical contributors and data producers from the International Cancer Genome Consortium (ICGC) for referencing the ICGC datasets.

